# The recruitment of indirect-waves within primary motor cortex during motor imagery: A directional Transcranial Magnetic Stimulation study

**DOI:** 10.1101/2022.05.10.491279

**Authors:** Cécilia Neige, Valentin Ciechelski, Florent Lebon

## Abstract

Motor imagery (MI) refers to the mental simulation of an action without any overt movement. While numerous transcranial magnetic stimulation (TMS) studies provided evidence for a modulation of corticospinal excitability and intracortical inhibition during MI, the neural signature within the primary motor cortex is not clearly established. In the current study, we used directional TMS to probe the modulation of the excitability of early and late indirect-waves (I-waves) generating pathways during MI. Corticospinal responses evoked by TMS with posterior-anterior (PA) and anterior-posterior (AP) current flow within primary motor cortex evoke preferentially early and late I-waves, respectively. Seventeen participants were instructed to stay at rest or to imagine isometric maximal contractions of the right flexor carpi radialis. We demonstrated that the increase of corticospinal excitability during MI is greater with PA than AP orientation. By using paired-pulse stimulations, we confirmed that short-interval intracortical inhibition (SICI) increased during MI in comparison to rest with PA orientation whereas we found that it decreased with AP orientation. Overall, these results indicate that the specific early I-waves generating pathway activated by PA orientation is probably more sensitive to the corticospinal excitability and intracortical inhibition modulations induced by MI.

## Introduction

Motor imagery (MI) is a cognitive process which refers to the mental simulation of an action without any overt movement (Jeannerod & Decety, 1995). MI is known to activate brain regions also involved during motor execution but is accompanied by a voluntary inhibition of the actual movement (Decety, 1996). Using vascular space occupancy method combined with high-resolution (7T) functional magnetic resonance imaging, Persichetti et al. (2020) supported the idea that MI activated only the superficial layers II/III with mainly cortico-cortical connections to the primary motor cortex (M1), whereas actual finger movements activated both superficial layers and the deeper layers Vb/VI with descending corticospinal projections. This would nicely explain the absence of muscle activity during MI. However, this exclusive activation of superficial layers within M1 during MI is at odds with numerous observations in the current literature.

Transcranial magnetic stimulation studies (TMS) provided evidence of the activation of the corticospinal pathway during MI, when compared to rest (Yahagi & Kasai, 1999; Grosprêtre *et al*., 2016). This activation is classically marked by an increase in the amplitude of the motor evoked potentials (MEPs) evoked by single-pulse TMS and recorded in the specific muscle involved in the imagined movement (Yahagi & Kasai, 1999; Lebon *et al*., 2012; Grosprêtre *et al*., 2016; Neige *et al*., 2020, 2021). According to Di Lazzaro and Ziemann (Di Lazzaro & Ziemann, 2013), the axons of the more superficial pyramidal neurons (P2/P3) are conceivably the most excitable neural elements to low-threshold TMS, due to their superficial location close to the stimulating coil. These axons also represent the main source of excitatory descending input to pyramidal tract neurons of layer V (Anderson *et al*., 2010). If TMS activates preferentially axons of superficial layer cells and MI induces an increase in TMS-evoked responses, it is most likely that superficial pyramidal neurons activated during MI directly excite deeper layers, contradicting the findings by Persichetti et al. (2020).

Previous studies that used TMS during MI rely on the interpretation of the MEPs amplitude evoked by the posterior-anterior (PA) current direction. However, MEPs amplitude is a complex and global readout that is thought to reflect the summation of several monosynaptic and polysynaptic descending inputs, termed D- (direct) and I-(indirect) waves, evidenced from spinal epidural recordings (Di Lazzaro *et al*., 2012). The first descending volley is thought to originate from direct activation (D-wave) of corticospinal tract axons, whereas the latter I-waves are thought to derive from indirect, trans-synaptic activation of the corticospinal neurons (Di Lazzaro & Rothwell, 2014a; Ziemann, 2020). These I-waves usually appear at ∼ 1.2–1.5 ms intervals are numbered in order of their appearance and are referred to as either early (I1) or late (I2, I3, I4) I-waves (Di Lazzaro *et al*., 2012; Ziemann, 2020). A non-invasively and valuable approach used to activate distinct sets of synaptic inputs to corticospinal neurons responsible for the early and late I-waves pathway is the directional TMS technique. It has been proposed that TMS-induced electric currents flowing from PA and AP (anterior to posterior) direction activate different sets of excitatory synaptic inputs that arrive at the pyramidal tract neurons and motor pathway several milliseconds apart (Di Lazzaro & Rothwell, 2014b; Di Lazzaro *et al*., 2017). PA stimulation preferentially elicits primarily early I-wave which is thought to originate from excitatory inputs to the basal dendrites of the corticospinal neurons in layer V of M1 (Di Lazzaro & Ziemann, 2013; Hannah, 2020). AP stimulation preferentially elicits later and more dispersed I-waves which are thought to result from mono-and poly-synaptic inputs from layers II/III of M1 (Ziemann, 2020), as well as the activation of horizontal cortico-cortical connections from surrounding brain regions to M1 (Di Lazzaro *et al*., 2017; Hannah, Cavanagh, *et al*., 2018). Therefore, the comparison between MEPs amplitude induced by PA and AP current directions allows us to infer about the different I-waves contributions evoked by separate subpopulations of interneurons.

In the current study, we investigated the modulation of early and late I-waves generating pathway specifically activated by PA and AP directed currents during MI and rest conditions. We used single-pulse and paired-pulse TMS to probe corticospinal excitability and short-interval intracortical inhibition (SICI), respectively. Interestingly, SICI affects mainly later I-waves that are mainly targeted by AP orientation (Nakamura *et al*., 1997; Hanajima *et al*., 1998; Cirillo & Byblow, 2016; Wessel *et al*., 2019), and SICI increases during MI, but only observed with PA orientation (Neige *et al*., 2020).

If MI preferentially recruits superficial layers within M1 (Persichetti *et al*., 2020), we would observe greater increase of corticospinal excitability and SICI with AP than PA orientation. On the contrary, if MI activates neural circuits downstream the pyramidal cells (Grosprêtre *et al*., 2015), we expect that MI would induce a specific modulation of the excitability of the early I-wave generating pathway activated by PA direction. This pathway would serve a critical role in modulating both corticospinal excitability and SICI.

## Material and Methods

### Participants

Seventeen healthy volunteers were recruited in the current study after providing written informed consent (3 females; age = 24.3 years, range 21-31 years; height = 177 ± 8 cm; weight = 69 ± 10 kg; right-handed as assessed by the Edinburgh Handedness Inventory (Oldfield, 1971)). All volunteers were screened by a medical doctor for contraindications to TMS (Rossi *et al*., 2009). The protocol was approved by the CPP SOOM III ethics committee (number 2017-A00064-49) and complied with the Declaration of Helsinki.

### Experimental setup

Participants were seated in an isokinetic dynamometer chair (Biodex System 3, Biodex Medical Systems Inc., Shirley, NY, USA). Participants’ right hand was firmly strapped in a neutral position to a custom-build accessory adapted for wrist movement recordings. The rotation axis of the dynamometer was aligned with the styloid process of the ulna. The upper arm was vertical along the trunk (shoulder abduction and elevation angles at 0°) and the forearm semipronated and flexed at 90°. First, participants familiarized themselves with the procedure of the voluntary force production feedback during an approximately 5-min warm-up of wrist flexions. They received visual feedback of the real time exerted force contraction on a computer screen located 1-m in front of them. Then, participants performed 3 maximal voluntary isometric contractions lasting 3 seconds with verbal encouragement, separated with at least 30 seconds rest in-between. The maximum of the three trials was defined as the participant’s maximal voluntary isometric contractions.

### Electromyographic recordings

Surface electromyographic (EMG) activity was recorded from the right flexor carpi radialis (FCR) using two silver-chloride (Ag/AgCl) electrodes placed over the muscle belly, at 1/3 of the distance from the medial epicondyle of the humerus to the radial styloid process. A ground electrode was placed over the medial epicondyle to the radial styloid. Signals were amplified (gain of 1000), band-pass filtered (10–1000 Hz), digitized at a sampling rate of 2000 Hz and stored for off-line analysis (Biopac Systems Inc. Goleta, CA, USA).

Background root mean square (RMS) of the surface EMG was calculated during the 100 ms epoch prior to TMS to ensure the absence of muscle contraction in each condition.

### Transcranial Magnetic Stimulation

Transcranial magnetic stimuli were applied using a 70-mm figure-of-eight coil through a Magstim BiStim² stimulator (The Magstim Co., Whitland, UK), with a monophasic current waveform. The optimal stimulation site on the scalp (hotspot) was defined as the location eliciting the largest MEP amplitude in the FCR muscle with PA-induced currents for a given intensity. For other coil orientations the same hotspot was used, since previous experiments have shown that the direction of the induced current does not significantly influence the position of the hotspot (Sakai *et al*., 1997; Hamada *et al*., 2013) (see Figure 1).

**Figure 1:**
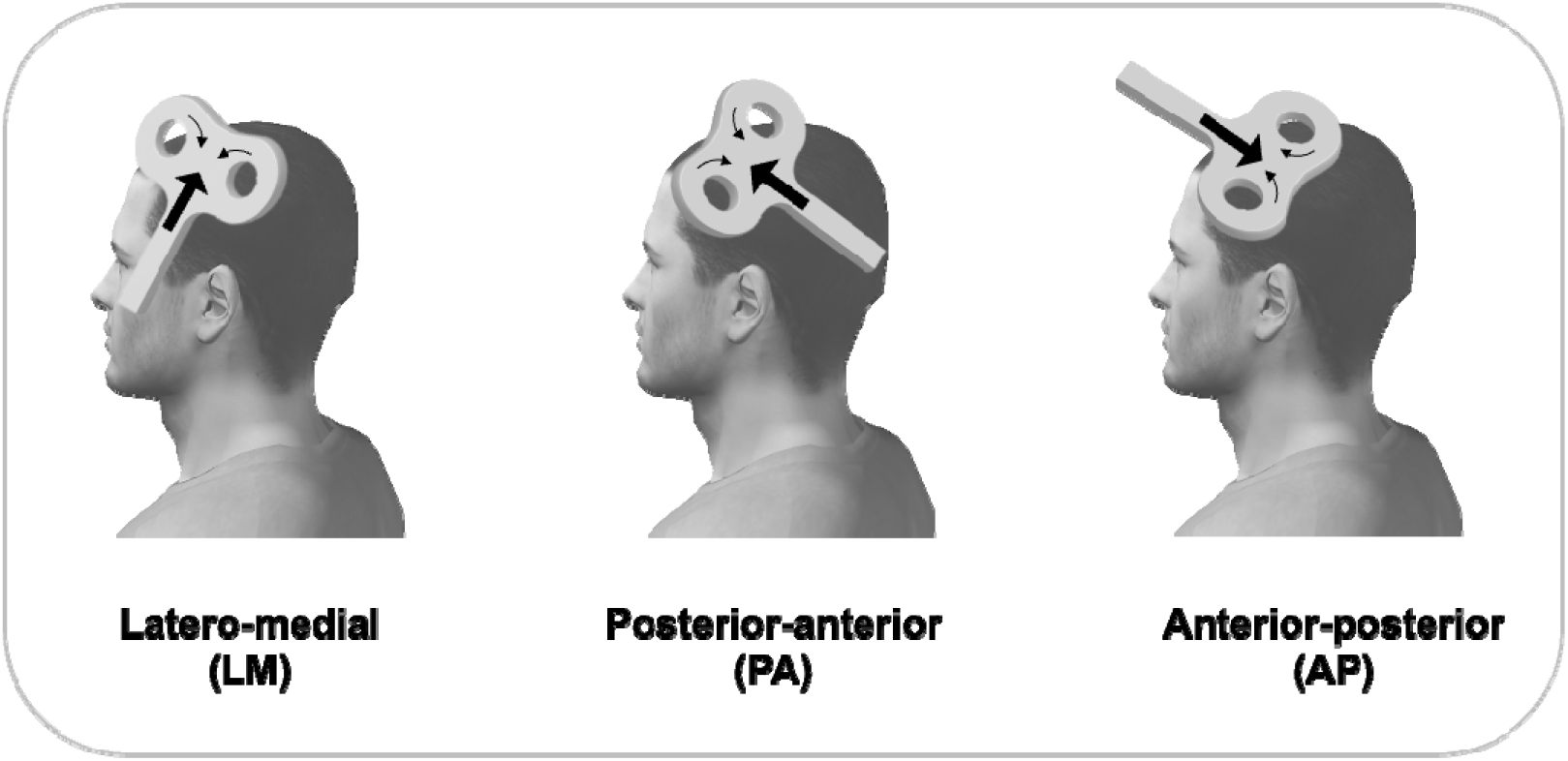
Illustration of the coil orientations and their direction of currents induced in the brain (large arrows) by single-and paired-pulse TMS.

The resting motor threshold (rMT) was determined for PA and AP directions as the lowest stimulus intensity required to evoke at least 5 MEPs of 50 _μ_V peak-to-peak amplitude out of 10 consecutive trials in the relaxed muscle (Rossini *et al*., 1994). The active motor threshold (aMT) was determined for PA, LM and AP directions as the lowest stimulus intensity required to evoke at least 5 MEPs of 200 _μ_V peak-to-peak amplitude out of 10 consecutive trials during 10% of the MVIC (Rossini *et al*., 1994).

### MEPs latency

The onset latency of MEPs obtained between PA-LM and AP-LM was used as an individual index of early (I1) and late (I2, I3) I-waves recruitment (Hamada *et al*., 2013; Neige & Beynel, 2020).

MEPs onset latency was determined for the FCR while participants maintained approximately 10% of their maximal voluntary isometric contractions (Hamada *et al*., 2013). This was done to ensure that low stimulus intensities could be used, thereby maximizing the selectively recruiting early or late I-waves with PA or AP currents. A higher stimulus intensity was used for LM to ensure that corticospinal neurons were directly stimulated (D-wave) at this coil orientation (Werhahn *et al*., 1994). Stimulation intensities was set at 110% of aMT_PA_, 110% of aMT_AP_ and 150% of aMT_LM_ (or 50% of maximum stimulator outputs (MSO) in participants whose 150% aMT_LM_ did not reach 50% MSO) (Hamada *et al*., 2013). Fifteen MEPs were recorded for each current direction, with the order of currents pseudo-randomized. The onset latency of MEPs assessed during muscle contraction was measured from the superimposed raw EMG waves-forms by visual inspection (Hamada *et al*., 2013; Hannah, Rocchi, *et al*., 2018).

### Adaptive threshold-hunting technique

To probe the distinct cortical elements recruited within M1 during MI, we used the adaptive threshold-hunting technique which consists in maintaining a constant MEP amplitude (called the MEP_target_, see below) by adjusting the TS stimulation intensity. The adaptive threshold-hunting paradigm offers several advantages when compared to the conventional protocol used to assess corticospinal excitability and SICI modulations. First, it allows to overcome the intrinsic MEPs amplitude variability thus providing more reliable results with shorter acquisition time (Samusyte *et al*., 2018). Then, it minimizes the potential “floor/ceiling effect” when complete inhibition is observed with the conventional SICI paradigm (Cirillo & Byblow, 2016). Finally, adaptive threshold-hunting technique relies on a weaker TS intensity (see below) than conventional paradigms (usually MEP _test_ 1mV or 120-130% rMT), which is thought to recruit more selectively early and later I-waves generating pathways (Di Lazzaro *et al*., 2001; Cirillo *et al*., 2020).

In the current study, unconditioned MEP_test_ and SICI modulation obtained during MI and compared to rest will be assessed by using the adaptive threshold-hunting technique with PA and AP current orientations known to preferentially elicit early-and late I-waves, respectively.

### MEP _target_

The hunting-threshold was defined as the TS intensity (expressed in percentage of the maximal stimulator output (%MSO)) required to elicit a MEP_target_ in the relaxed FCR muscle corresponding to the mean of 15 MEPs elicited at 115% rMT_PA_ in peak-to-peak amplitude. This led to a MEP_target_ of 0.251 ± 0.141 mV amplitude (see Table 1 for individual values). Generally, a non-personalized fixed 0.2 mV MEP_target_ amplitude is selected in studies using the adaptive threshold-hunting technique, corresponding approximately to 109% rMT (Fisher *et al*., 2002; Awiszus, 2003; Vucic *et al*., 2006; Menon *et al*., 2015; Cirillo & Byblow, 2016; Cirillo *et al*., 2018; Samusyte *et al*., 2018; Van den Bos *et al*., 2018; Neige *et al*., 2020). However, in the current study, a subject-specific MEP_target_ was chosen since 1) a huge between-subject variability in the intrinsic excitability of the corticospinal pathway exists, 2) a TS delivered at a lower intensity (i.e., below 110% rMT) could fail to evoke late I-waves, and limits SICI magnitude (Garry & Thomson, 2009) and 3) a TS delivered at a higher intensity can also elicit early I-waves when using an AP current direction, therefore limiting the interpretation differences obtained between PA and AP findings.

**Table 1:** Individual rMT and aMT (%MSO) according to PA and AP orientations. The individual MEP_target_ amplitude (mV) has been calculated from the mean of 15 MEPs elicited at 115% rMT_PA_.

| Subject | rMT<sub>PA</sub> | rMT<sub>AP</sub> | aMT<sub>PA</sub> | aMT<sub>AP</sub> | aMT<sub>LM</sub> | MEP<sub>target</sub> |
| --- | --- | --- | --- | --- | --- | --- |
| 1 | 40 | 46 | 26 | 35 | 30 | 0.213 |
| 2 | 32 | 41 | 28 | 35 | 29 | 0.384 |
| 3 | 50 | 35 | 28 | 39 | 35 | 0.109 |
| 4 | 46 | 55 | 39 | 48 | 40 | 0.095 |
| 5 | 39 | 51 | 37 | 44 | 44 | 0.079 |
| 6 | 32 | 32 | 25 | 28 | 29 | 0.337 |
| 7 | 45 | 49 | 39 | 41 | 44 | 0.291 |
| 8 | 45 | 56 | 35 | 50 | 35 | 0.647 |
| 9 | 43 | 55 | 33 | 47 | 34 | 0.255 |
| 10 | 30 | 35 | 26 | 30 | 31 | 0.234 |
| 11 | 50 | 52 | 43 | 46 | 43 | 0.170 |
| 12 | 36 | 40 | 27 | 34 | 30 | 0.162 |
| 13 | 35 | 48 | 29 | 42 | 33 | 0.258 |
| 14 | 38 | 42 | 31 | 36 | 34 | 0.368 |
| 15 | 43 | 49 | 34 | 42 | 37 | 0.263 |
| 16 | 40 | 45 | 36 | 41 | 39 | 0.313 |
| 17 | 41 | 49 | 32 | 42 | 35 | 0.119 |
| Mean | 40.3 | 45.9 | 32.2 | 40 | 35 | 0.252 |
| SD | 6.2 | 7.6 | 5.5 | 6.4 | 5.2 | 0.14 |

### Single-pulse TMS

The adaptive threshold-tracking single-pulse TMS technique was used to assess the unconditioned TS stimulation intensity required to reach the MEP_target_ amplitude (see General procedure) at rest vs. during MI, with PA and AP currents direction. The unconditioned TS intensity (expressed in %MSO) was quantified and compared across all experimental conditions. The more positive values characterize the higher TS Intensity required to reach the MEP_target_ amplitude.

### Short-interval intracortical inhibition (SICI)

The similar adaptive threshold-hunting technique with the same MEP_target_ amplitude was then used when conditioning the MEP_test_ with a sub-threshold conditioning TMS pulse (CS). This allows to assess SICI modulation at rest vs. during MI, with PA and AP currents direction. The CS intensity was fixed at 60% rMT_PA_ for SICI_PA_ and 60% rMT_AP_ for SICI_AP_, based on a previous study showing that higher CS intensities could lead to the unwanted recruitment of excitatory interneurons during MI, biasing the result interpretation (Neige *et al*., 2020). The inter-stimulus interval (ISI) between CS and TS was set at 3 ms, to induce the greatest inhibition when using AP current direction (Kujirai *et al*., 1993; Cirillo *et al*., 2018).

To probe the influence of the Task and Orientation on intracortical inhibition, the amount of SICI (expressed in INH%) was quantified for each condition using the following equation (Fisher *et al*., 2002):

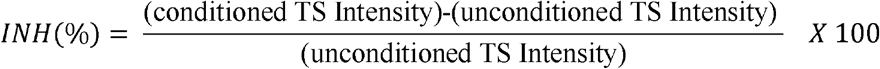

where positive values indicate inhibition and negative values indicate facilitation. The more positive values characterize the higher TS Intensity required to overcome the inhibitory influence of the CS and reach the MEP_target_ amplitude (Cirillo *et al*., 2020).

It has to be noted that only SICI data from 15 participants were used in the subsequent analysis because it was not possible to reach the MEP_target_ amplitude even at high stimulation intensity (>90% MSO) in 2 participants.

### General procedure

Experimental conditions (single-vs. paired-pulse; rest vs. MI; PA vs. AP current direction) were performed in different recording blocks and were randomized and counterbalanced across participants. An available online freeware (TMS Motor Threshold Assessment Tool, MTAT 2.0), based on a maximum-likelihood Parameter Estimation by Sequential Testing (PEST) strategy (Awiszus, 2003) was used with “assessment without a priori information” in line with previous studies (Cirillo & Byblow, 2016; Cirillo *et al*., 2018). The stimulation sequence always began with the TS at 37 %MSO. One experimenter held the coil over M1, while the other indicated whether (or not) the MEP amplitude was ≥ MEP_target._ The predictive algorithm then determined the next TS intensity to be delivered and was stopped after twenty stimulations, which provides sufficient accuracy for the threshold estimate according to previous studies (Awiszus, 2003, 2014; Ah Sen *et al*., 2017).

For MI trials, participants performed explicit and kinesthetic (somatosensory) MI of right wrist maximal isometric contractions with a first-person perspective for a duration of 3 s following an auditory cue (Hanakawa, 2016). The following instructions (in French) were carefully given to the participants: “When you hear the cue, try to imagine yourself performing the movement, to feel the movement, i.e., the muscle contraction and the tension that you would experience when performing the actual action. Be sure not to contract any muscles during the task and keep your eyes open” (Lebon *et al*., 2019; Neige *et al*., 2021). Kinesthetic MI strategy is thought to produce the greater muscle-specific and temporally modulated facilitation of the corticospinal pathway, compared to visual MI strategy (Stinear *et al*., 2006). The TMS pulses were triggered during the execution phase of MI trials and the intertrial interval was at least 4 s.

### Statistical analysis

Statistical analyses were performed using Statistical Program for the Social Sciences (SPSS) version 24 software (SPSS Inc., Chicago, IL, USA). Data distribution was assessed using the Shapiro-Wilk test. Homogeneity of variances was assessed by Mauchly’s test. If the sphericity assumption was violated a Greenhouse-Geiser correction was applied. Pre-planned post-hoc analyses were performed on significant interactions after applying a Bonferroni adjustment for multiple comparisons. Corrected p values for multiple comparisons are reported in the results section. The α level for all analyses was fixed at .05. Partial eta squared (η_p_^2^) values are reported to express the portion of the total variance attributable to the tested factor or interaction. For t-test analyses, effect sizes (Cohen’s d) are reported to indicate small (d = 0.2), moderate (d = 0.5) and large (d = 0.8) comparative effects. Values in parentheses in the text represent mean ± SD.

A first set of analyses were performed to control for potential methodological biases. A Student’s two-tailed paired sample *t-*tests were used to compare the rMT and aMT (%MSO) obtained for PA and AP current direction and to analyze the MEPs latency difference between PA-LM and AP-LM.

Then, two repeated-measures ANOVA were performed on the unconditioned TS Intensity (%MSO) and the SICI measurements (INH %) with two within-subject factors: Task_2_ (Rest vs. MI) and Orientation_2_ (PA vs. AP).

The RMS values were compared across conditions using a repeated-measures analysis of variance (ANOVA) with two within-subject factors: Task_2_ (Rest vs. MI), and Orientation_2_ (PA vs. AP). This ANOVA was performed separately for the unconditioned TS and the SICI measures. We predict no significant difference for these comparisons since an absence of any volitional muscle activity is expected for all the experimental conditions.

## Results

### Motor thresholds

Overall, both rMT (*t*(16) = −3.55, p = .003; Cohen’s *d* = -0.862) and aMT (*t*(16) = −8.147, p < .001; Cohen’s *d* = -1.976) were significantly smaller for PA compared to AP orientation (Table 1) as observed in previous studies using the adaptive-hunting threshold technique (Cirillo & Byblow, 2016; Cirillo *et al*., 2018).

### MEP latency

The analysis of MEPs latency difference revealed that PA-LM latency was significantly shorter compared with AP-LM latency (*t*(16) = -10.042, p < .001; Cohen’s *d* = -2.436). This result was consistent across participants (Figure 2) and suggested that the early wave recruited with PA orientation (mean latency = 17.06 ± 1.2 ms) and late I-waves recruited with AP orientation (mean latency = 18.72 ± 1.4 ms) could be differentially recruited within individuals.

**Figure 2:**
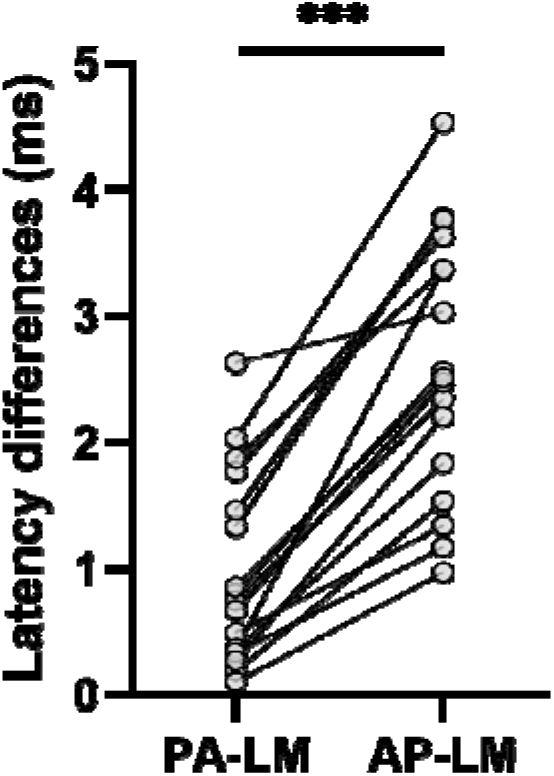
Individual MEPs onset latency differences (in ms) obtained between PA-LM and AP-LM current orientations. PA: posterior-anterior; LM: latero-medial; AP: anterior-posterior.

### Unconditioned TS Intensity

Figure 3A illustrates the unconditioned TS Intensity obtained at rest and during MI for PA and AP current directions. We found a significant main effect of Orientation (F_(1,16)_ = 39.338, p < .001; n_p_^2^ = .711) indicating that the unconditioned TS Intensity required to reach the MEP_target_ was significantly higher for the AP orientation than the PA orientation. A main effect of Task was also observed (F_(1,16)_ = 11.004, p = .004; n_p_^2^ = .409) but more importantly the Orientation by Task interaction was significant (F_(1,16)_ = 5.130, p = .038; n_p_^2^ = .243). Post hoc analyses revealed that for both PA and AP orientations, the unconditioned TS Intensity required to reach the MEP_target_ was significantly lower during MI than at rest (p =.005 for PA and p =.025 for AP) indicating that MI increased corticospinal excitability. Importantly, when comparing the rest vs. MI ratios according to the Orientation, the reduction of the unconditioned TS Intensity for MI when compared to rest was significantly more important for PA than AP direction (*t*(16) = -2.601, p = .019; Cohen’s *d* = -0.631) (see Figure 3B). This suggests that the classical corticospinal excitability increase during MI is mainly driven by early I-waves recruitment.

**Figure 3:**
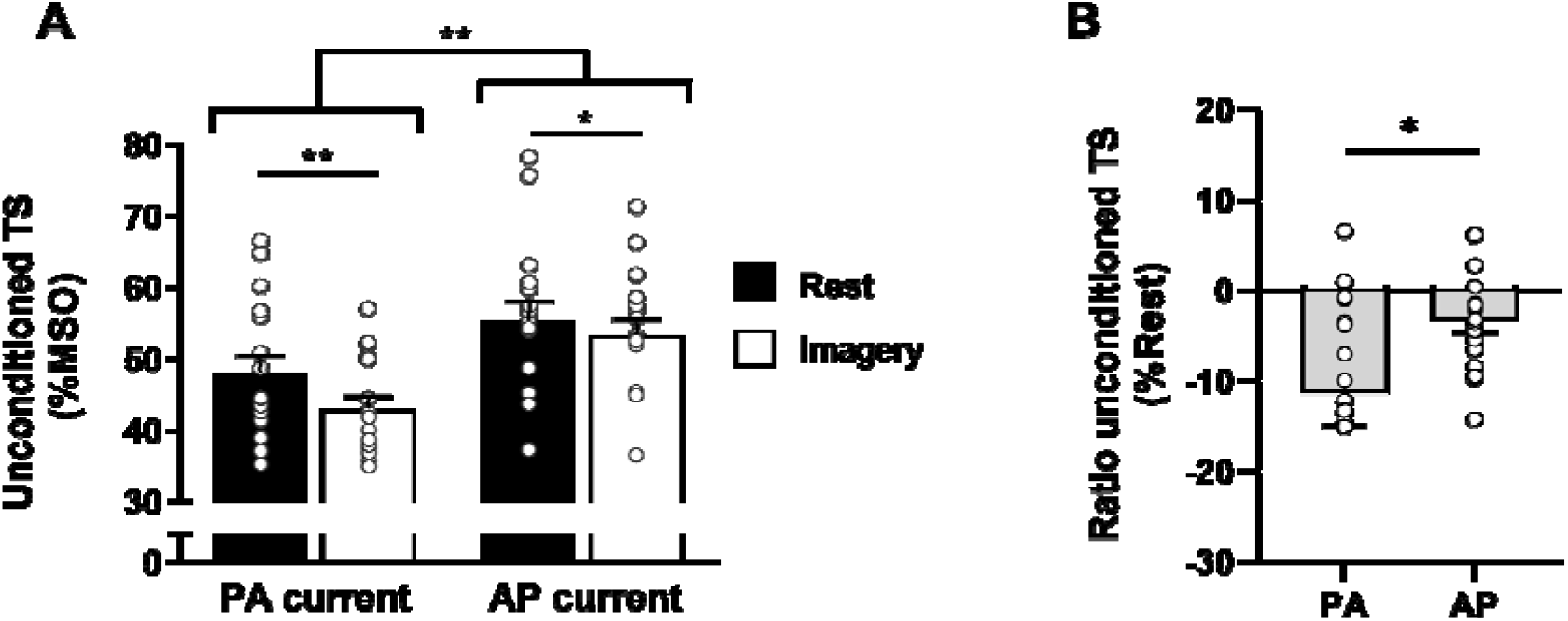
**A)** Mean ± SE for the unconditioned TS Intensity (%MSO) obtained with the hunting-threshold technique at rest and during motor imagery for the two current orientations. Lower values of %MSO indicate lower TS intensities to reach the MEP_target_ amplitude. **B)** Ratio for the unconditioned TS Intensity obtained during motor imagery and expressed as a percentage of rest condition for the two current orientations. Negative values indicate lower TS intensities during Imagery in comparison to rest, and therefore an increase of corticospinal excitability, which is greater with PA orientation than with AP orientation. Data points represent individual participants. PA: posterior-anterior; AP: anterior-posterior. *p< .05; **p < .01.

### Conditioned TS Intensity (SICI)

Figure 4 illustrates the percentage of inhibition (SICI) obtained at rest and during MI for the PA and AP current direction. We did not find any main effects of Orientation (F_(1,14)_ = 1.107, p = .311) or Task (F_(1,14)_ < 1, p = .895), but the Orientation by Task interaction was significant (F_(1,14)_ = 11.995, p = 0.004, n_p_^2^ = .461). Post hoc comparisons showed that at rest, the amount of SICI was higher for AP orientation than the PA orientation (p = .031) whereas it was not significant when comparing orientations during MI (p = .106). Moreover, SICI was greater during MI compared to rest with the PA orientation (p = .028), whereas SICI was lower during MI compared to rest with the AP orientation (p = .033).

**Figure 4:**
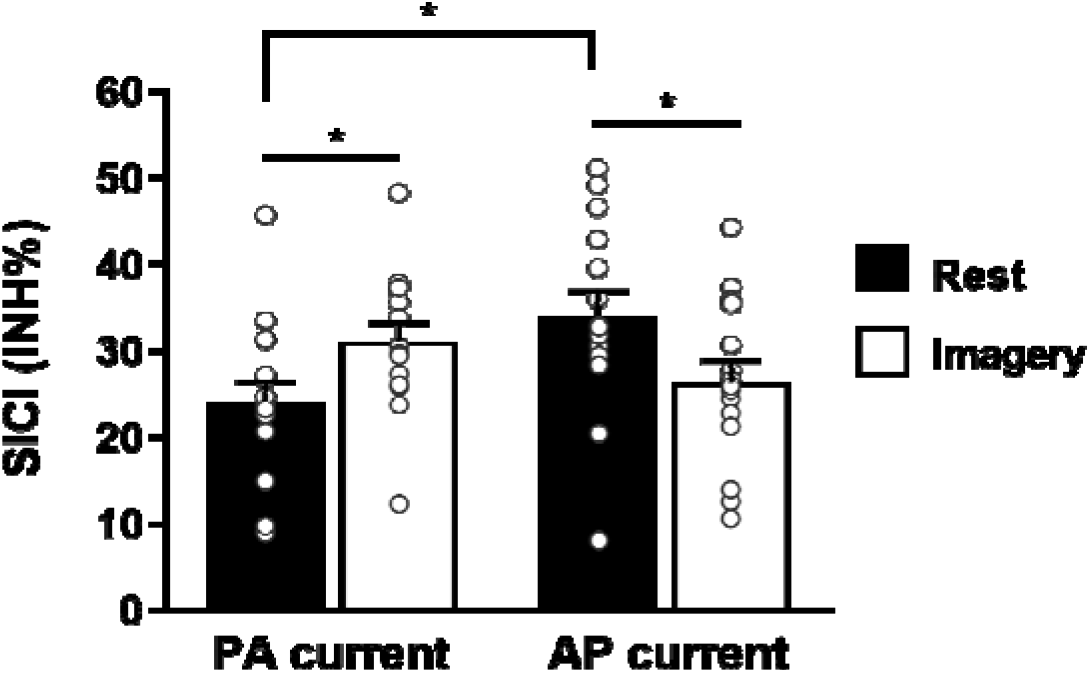
Mean ± SE for the SICI (%INH) obtained with the hunting-threshold technique at rest and during motor imagery for the two current orientations. Data points represent individual participants. PA: posterior-anterior; AP: anterior-posterior. *p < .05

### RMS

The analysis of RMS of EMG background for the unconditioned TS showed no significant difference between PA and AP orientation (F_(1,16)_ < 1, p = .548) and more importantly between rest and MI (F_(1,16)_ < 1, p = .542). Nor was the Orientation by task interaction (F_(1,16)_ = 3.594, p = .076).

Similarly, the analysis of the RMS values for the conditioned TS yielded no significant main effect of Orientation (F_(1,14)_ < 1, p = .997), Task (F_(1,14)_ = 1.77, p = .204) or Orientation by Task Interaction (F_(1,13)_ < 1, p = .563).

Together, these results indicate that any changes in corticospinal excitability cannot be attributed to differences in the EMG levels prior to the TMS pulse.

## Discussion

In the current study, we demonstrated for the first time that MI activates different subsets of neurons within M1, by means of directional TMS and the adaptive threshold-hunting technique. The increase of corticospinal excitability during MI may originate from an increase in the excitability of early I-waves rather than late I-waves as evidenced by a greater increase observed with PA orientation when compared to AP orientation. By using paired-pulse stimulation, the results confirmed that the amount of SICI measured at rest is higher for AP orientation than the PA orientation (Neige *et al*., 2020). Interestingly, SICI increases observed during MI (vs. at rest) was also restricted to PA orientation. On the contrary, when using AP orientation SICI was lower during MI compared to rest. Taken together, it suggests that the specific later I-waves evoked by AP orientation is less sensitive to the corticospinal excitability and intracortical inhibition modulations induced by MI. This result confirms the hypothesis that MI activates neural circuits downstream the pyramidal cells and produces a subliminal motor output that reaches the spinal level (Grosprêtre *et al*., 2015) rather than induce a specific superficial activation restricted to the superficial layers within M1 that could explain the absence of muscle activity during motor imagery (Persichetti *et al*., 2020). The excitability of the early I-wave generating pathway activated by PA orientation may serve a critical role in modulating both corticospinal excitability and intracortical inhibition.

### Corticospinal excitability increases observed during MI is greater for PA than AP orientation

It is well known that MEPs amplitude increases during MI compared to rest reflects an increase of neuron responsiveness to TMS (Grosprêtre *et al*., 2016). However, MI is a complex state that also involves mechanisms that actively suppress the transmission of the motor command into the efferent pathway, supporting the action of inhibitory pathways during MI (Jeannerod & Decety, 1995). Actually, it is still unclear which inhibitory mechanisms counteract the corticospinal excitability increase in order to prevent the production of an overt movement.

Based on previous reports demonstrated that single-pulse TMS with a PA orientation preferentially recruits early I-waves (I1), whereas AP orientation preferentially recruits later I-waves (I3) (Zoghi *et al*., 2003; Di Lazzaro & Rothwell, 2014b; Di Lazzaro *et al*., 2017), we used directional TMS to activate different sets of excitatory synaptic inputs within M1.

First, our results demonstrate that our experimental setting (coil orientation) was correct, with a significant latency difference between PA-LM and AP-LM, supporting a differential recruitment of cortical neurons in M1 relative to the current orientation (Werhahn *et al*., 1994; Hamada *et al*., 2013; Di Lazzaro & Rothwell, 2014). If it has already been shown in a forearm extensor muscle (McCambridge *et al*., 2015), our study is the first to demonstrate latency difference in the FCR muscle.

In an attempt to further investigate the involvement of different subsets of cortical neurons during MI compared to resting state, we used the adaptive threshold-hunting technique and compared the unconditioned TS intensity (%MSO) required to reach the MEP_target_ at rest and during MI, with PA and AP orientation. We found that the unconditioned TS Intensity required to reach the MEP_target_ was significantly lower during MI than at rest for both the PA and the AP orientations. However, when comparing the ratios MI/rest according to the orientation, the reduction of the unconditioned TS Intensity for MI was significantly more important for PA than AP direction. Taken together, these results indicate that early I-wave generating pathway within M1 possibly mediated the MEPs amplitude increases observed during MI. The exact underpinning neurophysiology of I-waves generation remains largely misunderstood (Ziemann, 2020). However, it has been suggested that the early I-wave evoked by TMS with PA orientation is the result of the activation of monosynaptic cortico-cortical connections projecting onto the large corticospinal neurons of layer V (Di Lazzaro & Ziemann, 2013; Di Lazzaro *et al*., 2017; Hannah, 2020). This result seems coherent with previous literature assuming that the early I-wave is produced by a different anatomical substrate and mechanism than the late I-waves (Ziemann, 2020). Crucially, specific early I-wave evoked by TMS with PA orientation is thought to be enhanced by increases in corticospinal excitability (Di Lazzaro *et al*., 1998a, 2017), which was the case during MI. For example, voluntary muscle contraction increased both corticospinal excitability and the relative contribution of the early I-wave (Di Lazzaro *et al*., 1998a, 2017).

The findings of the current study also extend and consolidate our knowledge regarding the distinct I-wave circuits recruitment during behavioral states that share analogous control mechanisms and neural circuits with overt movements but without any muscle activity (i.e., covert actions) (Hannah, 2020). Indeed, recent studies exploited the directional TMS technique to probe the different subset of cortical neurons recruited during motor preparation (Hannah, Cavanagh, *et al*., 2018) and action observation (Hannah, Rocchi, *et al*., 2018). The results showed that during motor preparation, the decrease of the corticospinal excitability in the selected and non-selected muscles was accompanied by a selective suppression of the subset of excitatory inputs to corticospinal neurons responsible for late I-waves, while the subset responsible for early-I wave remains unaffected (Derosiere, 2018; Hannah, Cavanagh, *et al*., 2018). On the contrary, when using directional TMS technique during action observation, other authors failed to observe a selective recruitment of the early or late I-waves pathway, probably due to a large intersubject variability in corticospinal modulations (Hannah, Rocchi, *et al*., 2018). Overall, the recent use of directional TMS technique applied during motor preparation, action observation and during MI allow to gain further insight into the distinct circuitry recruited with TMS that contributes to the corticospinal excitability modulation.

### SICI increases during MI is restricted to PA orientation

The adaptive threshold-hunting paired-pulse TMS technique was also used in the current study to examine modulations of SICI during MI and at rest and how they are influenced by TMS coil orientation.

SICI involves a subthreshold CS which is thought to activate low-threshold inhibitory interneurons that employ-aminobutyric acid type A receptor (GABA_A_). The effect of the activation of these GABAergic inhibitory interneurons is the reduction of the excitatory inputs activated by the TS (Kujirai *et al*., 1993; Di Lazzaro *et al*., 1998b, 2017). Importantly, it has been described previously that SICI affects predominantly later I-waves (I3) which are mainly targeted by AP orientations (Nakamura *et al*., 1997; Hanajima *et al*., 1998; Di Lazzaro *et al*., 2012; Higashihara *et al*., 2020). Moreover, it is known that the use of the adaptive threshold-tracking technique with an AP induced current with a 3-ms ISI provides a more robust and sensitive measure of SICI than with a PA induced current (Cirillo & Byblow, 2016). Therefore, a greater level of SICI assessed at rest using AP-compared with PA-orientation demonstrated in the present study corroborates and replicates earlier findings (Cirillo & Byblow, 2016; Cirillo *et al*., 2018, 2020). This result also demonstrates further evidence that SICI is mediated by the recruitment of inhibitory interneurons generating late I-waves.

By comparing the extent of SICI modulation obtained with a PA induced current, we found that there was significantly more inhibition during MI when compared to rest. This finding also corroborates a previous study showing that when tested with low CS intensities, as in the current study (< 70 %rMT), SICI is greater during MI than at rest (Neige *et al*., 2020). This increase of SICI could reflect the crucial role played by cortical interneurons within M1 in the fine-tuning neural processes required during MI. This may prevent the production of an overt movement when the mental representation of that movement is activated.

Conversely, by comparing the extent of SICI modulation obtained with an AP induced current, we found a SICI decrease during MI compared to rest. Moreover, contrary to what was found during the resting state, the level of SICI assessed during MI using AP-compared with PA-orientation was not significantly greater. These results, combined with the unconditioned TS intensity findings indicate that the specific later I-waves evoked by AP orientation are probably less sensitive to the corticospinal excitability and intracortical inhibition modulations induced by MI.

### MI influences a specific distributed circuit that can differentially contribute to early and late I-waves

Neuroimaging studies provided evidence that MI activates a premotor-parietal network including cortical and subcortical brain regions such as dorsolateral prefrontal cortex, supplementary motor area, premotor cortex, posterior parietal regions, putamen and cerebellum (Hardwick et al. 2018). Crucially, M1 is known to integrate inputs from some of these structures and the latter are differentially recruited according to current orientation. For example, late I-waves evoked by AP orientation could activate axons of neurons of the premotor cortex projecting to the corticospinal cells (Groppa *et al*., 2012; Volz *et al*., 2015; Aberra *et al*., 2020; Siebner, 2020; Desmons *et al*., 2021). Recently, Oldrati et al. (2021) reported that following offline 1Hz inhibitory repetitive TMS over the dorsal premotor cortex (PMd), corticospinal excitability assessed during kinesthetic MI was not significantly higher than rest condition (Oldrati *et al*., 2021). These findings suggest a facilitatory connectivity from PMd to M1 during MI. Although this remains speculative, it is possible that the facilitatory input from PMd to M1 occurring during MI has decreased the SICI level within M1. Moreover, the opposite higher level of SICI during MI (vs. rest) observed with PA current reflect the activation of inhibitory inputs received from somatosensory cortex and SMA, both areas known to functionally inhibit M1 when imagining (Kasess *et al*., 2008; Oldrati *et al*., 2021). Finally, it is also possible that cerebellum, which facilitates M1 excitability during MI (Tanaka *et al*., 2018; Monany *et al*., 2021), also contributes to the result of the current study since the influence of cerebellum on M1 might occur via interactions with specific I-waves generating circuits (Spampinato *et al*., 2020).

### Limitations and perspectives

Several limitations need to be taken into consideration when interpreting the results of this study. First, SICI modulations tested with the adaptive-threshold hunting technique also depend on CS intensity (particularly during MI) (Vucic *et al*., 2009; Ibáñez *et al*., 2020; Neige *et al*., 2020) and interstimulus-intervals (Fisher *et al*., 2002). These two parameters were not manipulated in the current study and the careful consideration of stimulation parameters selected for SICI assessment deserve further investigations. Moreover, the activation of distinct subsets of neurons within M1 according to the PA or AP orientation is known to be sensitive to specific stimulation parameters such as pulse duration, pulse shape and phase amplitude (D’Ostilio *et al*., 2016; Hannah & Rothwell, 2017; Hannah *et al*., 2020; Spampinato, 2020). Due to technical limitations, these parameters were not optimized to selectively activate inputs to corticospinal neurons with PA and AP orientations. Future studies should consider this limitation to enhance the efficiency of the directional TMS technique to probe distinct PA and AP sensitive neurons during MI.

In order to gain further insight into the different subsets of cortical neurons and interneuronal circuits recruited during MI, it would be worthwhile to exploit recent techniques also developed to probe the separate subsets of inputs in M1. For example, Kurz et al. (2019) developed a novel non-invasive method that combines single-pulses TMS with peripheral nerve stimulations of the median nerve generating an H-reflex. This technique makes it possible to estimate excitability changes of different micro-circuits of M1 which reflect layer-specific activity (Dukkipati & Trevarrow, 2019; Kurz *et al*., 2019). Since layer-specific cortical circuits activity has been recently evidenced during MI (Persichetti *et al*., 2020) and that corticospinal neurons responsible for the early and late I-waves pathways are thought to originate from layer-specific cortical circuits, the technique of Kurz et al. could be a promising tool to delineate further the different subsets of neurons in M1 activated during MI. Finally, the exact contribution of the early and late I-waves can be captured by delivering paired-pulse TMS at precise intervals approximating the different I-waves latency (Tokimura *et al*., 1996; Hanajima *et al*., 2002). This technique has been recently applied during grasping observation to isolate the contribution to early and late excitatory inputs to M1 (Cretu *et al*., 2020) and could be tested during MI.

In conclusion, this study is the first to present evidence that the increase of corticospinal excitability and intracortical inhibition during MI may originate from a specific modulation of the excitability of the early I-wave (rather than later I-waves) generating pathway. This finding is reflected by a greater corticospinal excitability increase observed during MI (compared to rest) with PA than AP orientation. Moreover, the SICI increase during MI was only restricted to PA orientation. We found decreased SICI when using AP orientation, which is more sensitive to later I-waves generating pathway. Taken together, the results confirm that MI modulates preferentially the excitability of the early I-wave generating pathway.

## Acknowledgments

This research was financially supported by the ‘Investissements d’Avenir’ French program, project ISITE-BFC (contract ANR-15-IDEX-0003).

CN was involved in conception and study design, acquisition of data, data/statistical analysis with interpretation, manuscript preparation, and final approval of the manuscript.

VC was involved in acquisition of data, data analysis and final approval of the manuscript.

FL was involved in conception and study design, data interpretation, supervision, funding, manuscript preparation and final approval of the manuscript.

The authors want to thank Dr. Anaïs Gouteron for participant’s medical inclusion, William Dupont and Dylan Rannaud Monany for technical assistance during acquisition of data.

